# Dual regulation of Misshapen by Tao and Rap2l promotes collective cell migration

**DOI:** 10.1101/2023.07.21.550060

**Authors:** Gabriela Molinari Roberto, Alison Boutet, Sarah Keil, Gregory Emery

## Abstract

Collective cell migration occurs in various biological processes such as development, wound healing and metastasis. During Drosophila oogenesis, border cells (BC) form a cluster that migrates collectively inside the egg chamber. The Ste20-like kinase Misshapen (Msn) is a key regulator of BC migration coordinating the restriction of protrusion formation and contractile forces within the cluster. Here, we demonstrate that the kinase Tao acts as an upstream activator of Msn in BCs. Depletion of Tao significantly impedes BC migration and produces a phenotype similar to Msn loss-of-function. Furthermore, we show that the localization of Msn relies on its CNH domain, which interacts with the small GTPase Rap2l. Our findings indicate that Rap2l promotes the trafficking of Msn to the endolysosomal pathway. When Rap2l is depleted, the levels of Msn increase in the cytoplasm and at cell-cell junctions between BCs. Overall, our data suggest that Rap2l ensures that the levels of Msn are higher at the periphery of the cluster through the targeting of Msn to the degradative pathway. Together, we identified two distinct regulatory mechanisms that ensure the appropriate distribution and activation of Msn in BCs.

## INTRODUCTION

Collective cell migration is involved in various biological processes, including normal development, tissue homeostasis and cancer metastasis. Cells can migrate collectively as cell sheets, streams, or clusters [1, 2]. Recent studies have revealed that cancer cells migrating as clusters are more efficient at forming metastases compared to single cells [3–5]. Consequently, it is imperative to gain a deeper understanding of the regulatory mechanisms that govern the ability of cells to migrate collectively.

Border cells (BCs) in the Drosophila ovary is a robust model for studying the collective cell migration of clusters in vivo. The ovary contains egg chambers that consist of 15 nurse cells and one oocyte, surrounded by somatic epithelial follicular cells. During stage 9 of oogenesis, a subset of follicular cells at the anterior end of the egg chamber differentiates into BCs, forming a cluster of 6 to 10 cells. This cluster detaches from the follicular epithelium and undertakes an invasive migration, moving through the nurse cells toward the oocyte [6, 7].

The migration of BCs is guided by ligands secreted by the oocyte which activate receptor tyrosine kinases [8–10]. Subsequently, this activation leads to the activation of the small GTPase Rac and the formation of protrusions [11]. Interestingly, Rac activity and protrusion formation are restricted to one or two cells at the front of the cluster that become the leader cells [12]. Recent research has delved into the mechanisms that restrict protrusion formation to leading cells. Several studies have shown that the presence of a supracellular actomyosin network at the periphery of the cluster prevents the formation of protrusions in non-leader cells. The assembly of this supracellular network requires the plasma membrane and actin-binding protein, Moesin (Moe) [13–17]. Knockdown of Moe or Myosin II induces ectopic protrusions in non-leader cells and blocks migration [13, 15, 17]. Furthermore, BC migration requires Myosin II-mediated contractility for the detachment of the cluster from the rest of the follicular epithelium and the retraction of protrusions [14, 18-20].

Recently, we identified the Ste20-like kinase Misshapen (Msn) as a key regulator of BC migration coordinating protrusion formation with contractile forces [14]. We showed that Msn regulates protrusion restriction to the leader cells by directly phosphorylating Moe at the periphery of the cluster. Moreover, Msn promotes Myosin II-mediated contractility through a Moe-independent mechanism regulating protrusions dynamics and detachment of the cluster. As such, Msn activity at the periphery of the BC cluster is essential for coordinated migration. Therefore, we rationalized that the mechanisms that control the spatiotemporal activity of Msn may play an important role in BCs.

Msn is composed of a kinase domain and a Citron homology (CNH) domain separated by a long coiled-coil region (Fig.1A). Activation of Msn requires the phosphorylation of its kinase domain. In Drosophila midgut enteroblasts, the kinase Tao phosphorylates the conserved threonine residue (T194) at the Msn kinase domain, promoting its activation [21]. However, in BCs, in addition to the kinase activation, Msn function requires its proper localization at the cell cortex to phosphorylate Moe [14]. Yet, Msn is devoid of known membrane localization domain, suggesting that its localization at the cell cortex depends on its interactions with other proteins. Interestingly, the CNH domain of different proteins can bind to Rho and Rap small GTPases [22, 23] and it was shown to regulate the subcellular localization of Msn orthologues [24, 25].

**Figure 1:**
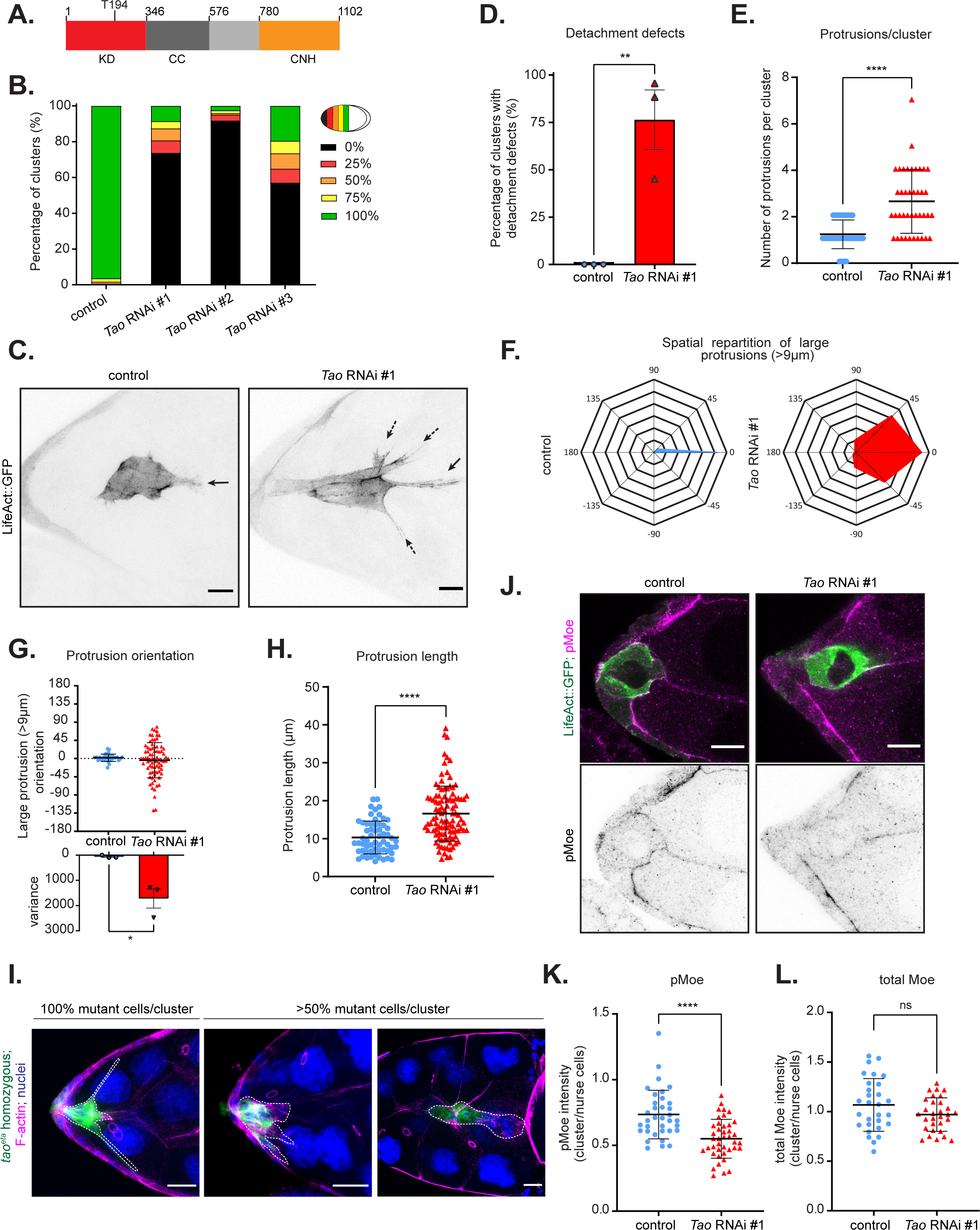
Tao is required for protrusion restriction and cluster detachment (A) Illustration of Msn protein structure with its different domains and the Tao phosphorylation site. KD - kinase domain, CC-coiled-coil and CNH - CNH domain. (B) Quantification of the percentage of BC clusters at each sector in stage 10 egg chambers after expression of a control RNAi or three independent dsRNA against *Tao*. (n= 306, 373, 121 and 128, respective to the histogram order, N=3) (C) Representative Z-projection of LifeAct::GFP in control and Tao-depleted clusters shown in inverted greyscale. Plain arrows indicate protrusions at the front of the clusters, and dotted arrows indicate protrusions at the side. Scale bar: 10µm (D) Quantification of the percentage of clusters that have not detached from the follicular epithelium by stage 10 (fixed samples) (n>60; N=3 ;**: p<0.01, unpaired Student’s t-test). (E) Quantification of the number of protrusions per cluster in control and Tao-depleted clusters. (n = 51 and n = 40, respectively, N=3, ****: p<0.0001, Mann-Whitney test). Any protrusions longer than 4 μm were manually counted using fixed samples expressing LifeAct::GFP. (F) Analysis of the orientation of protrusions longer than 9μm in control and Tao-depleted clusters. Grid space scale correspond to 5 protrusions (n = 51 and n = 40, respectively, N=3) (G) Quantification of the angle between the tip of protrusions longer than 9μm and the direction of migration in control and Tao-depleted clusters. The variance between experiments is shown in the graph below (n = 51 and n = 40, respectively; N=3, *: p<0.05, unpaired Student’s t-test) (H) Quantification of the length of protrusion in control and Tao-depleted clusters. (n = 60 and 103, respectively; N=3, ****: p<0.0001, Mann-Whitney test). (I) Representative images of complete and mosaic homozygous *tao^eta^* mutant BC clusters. Homozygous *tao^eta^* mutant clones express GFP and a white dashed line contours the BC cluster. Scale bar: 10µm. (J) Representative images of the pMoe immunostainings in control and Tao-depleted clusters at the onset of migration. The pMoe channel is displayed individually as inverted greyscale images. Scale bar: 10µm. (K) Quantification of the ratio of pMoe mean intensity at the cluster periphery, normalized to the signal between nurse cells after expression of control RNAi and *Tao* RNAi. (n= 35 and 41, respectively, N=3, ****: p<0.0001, Mann-Whitney test) (L) Quantification of the ratio of total Moe mean intensity at the cluster periphery, normalized to the signal between nurse cells, after expression of control RNAi and *Tao* RNAi. (n= 29 and 31, respectively, N=3, ns: not significant, Mann-Whitney test). All error bars represent SEM.

Here, we identified that Tao is required for BC migration by activating Msn. Depletion of Tao impairs BC migration and produces a loss-of-function phenotype similar to Msn. However, it does not affect Msn localization. We show that the localization of Msn at the cell cortex depends on its CNH domain which interacts with the small GTPase Rap2l. Accordingly, we found that Rap2l is required for BC migration and controls the balance of Msn levels at the different cell-cell junctions of BC clusters by promoting the trafficking of Msn to late endosomes. Depletion of Rap2l reduces Msn levels in late endosomes and increases the overall levels of Msn in particular at the cell-cell junctions between BCs. Overall, our research reveals a dual regulation of Msn: its distribution along different junctions of the BC cluster is regulated through a Rap2l-mediated degradation while its activity at the periphery of the cluster is promoted by its phosphorylation by Tao.

## RESULTS

### Tao depletion phenocopies Msn loss-of-function in BCs

The recent identification of Tao as an upstream kinase of Msn in Drosophila midgut enteroblasts [21] led us to speculate whether Tao could regulate Msn during BC migration. The loss of Msn in BCs has been associated with two distinct phenotypes. First, it results in contractility defects, which cause elongated protrusions and impede cluster detachment. Second, it leads to the emergence of multiple and misorientated protrusions [14]. Here, we hypothesize that if Tao is the upstream kinase of Msn in BCs, clusters depleted of Tao should exhibit a phenotype that closely resembles the loss-of-function phenotype of Msn.

To test this hypothesis, we first determined whether Tao is required for collective cell migration. We used the BC specific *c306*-Gal4 driver to express three independent double-stranded RNAs (dsRNAs) targeting non-overlapping sequences of *Tao*. The three different RNAi lines significantly reduce protein levels of Tao (Fig.S1A,B) and produce similar phenotypes (Fig.1B-H, Fig.S1C-H). These RNAi lines were previously used to characterize the role of Tao [21, 26-31]. We found that while control BC clusters completed their migration and were located next to the oocyte in stage 10 (S10) egg chambers, less than 20% of the clusters depleted of Tao could complete the migration on time. In fact, most clusters expressing *Tao* RNAi were unable to complete the first 25% of the migration path and remained at the onset of migration (Fig.1B). These data indicate that Tao is required for normal BC migration.

To further investigate the function of Tao in collective cell migration, we conducted a phenotypic analysis of BC clusters depleted of Tao. To assess the morphology of the cluster, we express the actin probe LifeAct fused to GFP (LifeAct::GFP) in BCs. Unlike control clusters, we observed that Tao-depleted clusters remained attached to the follicular epithelium in the S10 egg chamber (Fig.1C,D). Moreover, at the onset of migration, we noted a significant increase in the number of protrusions per cluster when Tao is depleted compared to the control (Fig.1C, E). For consistency, the rest of our phenotypically analysis was performed on clusters at the onset of migration, as it is the moment when the migration of most clusters is impaired, unless specifically mentioned.

To quantify the spatial distribution of protrusions, we measured the angle between large protrusions (longer than 9µm) and the migration direction. Furthermore, we represented the position of protrusions using a radar map, as previously [15]. At the onset of migration, control clusters have most of their protrusions aligned towards the migration direction. However, in Tao-depleted clusters, we observed a lack of this directional alignment and instead noticed a more scattered orientation of protrusions (Fig.1F-G). Furthermore, clusters expressing *Tao* RNAi exhibited longer protrusions than control clusters (Fig.1H). Our results suggest that Tao plays a critical role in cluster detachment and regulates the restriction of protrusions to the leader cells.

To further confirm the loss-of-function phenotype observed, we generated clones with the null allele *tao^eta^* [32]. We used the MARCM technique [33] to identify homozygous mutants BCs by the expression of GFP. Only six well-formed clusters in S10 egg chambers were found to be composed entirely of mutant cells. Notably, none of these clusters detached from the follicular epithelium. Furthermore, we noticed a significant delay in the migration of mosaic clusters in which more than 50% of cells were mutants. Interestingly, mutant cells were consistently positioned at the rear of the cluster, with many of them remaining attached to the follicular epithelium (Fig.1I). We also observed significant morphological changes in the mutant clusters as they present misorientated and longer protrusions (Fig.1I). Altogether, our data show that Tao is required to restrict protrusions and promotes detachment of BC clusters.

Msn regulates restriction of protrusion by phosphorylating Moe at the periphery of the cluster. Accordingly, depletion of Msn reduces the levels of phosphorylated Moe (pMoe) [14]. To determine if a similar mechanism is at play in Tao-depleted clusters, we measured the levels of pMoe. We observed a significant decrease in pMoe levels at the periphery of clusters expressing *Tao* RNAi. (Fig.1J-K, Fig.S1G). In contrast, total Moe levels were unaffected (Fig.1L, Fig.S1H). These results suggest that Tao regulates the restriction of protrusion to the leader cells through Moe phosphorylation.

### Tao activates Msn but does not affect its localization

The similarities between Msn and Tao loss-of-function phenotypes support the hypothesis that Tao is an upstream kinase of Msn in BCs. To further investigate this relationship, we examined both Msn and Tao localization and compared their distribution. Using an antibody against Msn, we observed that Msn is localized at the cell cortex and, to a lesser extent in the cytoplasm. During onset of migration, Msn is enriched at the inner junctions (the interface between BCs) and at the periphery of the cluster (the interface between BCs and nurse cells). Its distribution mid-migration reveals a preference for the cluster periphery, as previously described [14] (Fig.2A). These findings were confirmed with a knock-in GFP line of Msn (GFPki-Msn), which endogenously expresses Msn with a GFP tag [21] (Fig. S2A,B). As the anti-Msn antibody was not previously described, we validated its ability to label Msn by staining clusters expressing GFPki-Msn. As expected, the pattern revealed by the antibody is almost identical to GFPki-Msn (Fig.S2C). Furthermore, depletion of Msn in cluster led to a strong decrease of the signal (Fig.S2D). To determine if Tao colocalizes with Msn, we employed a knock-in line, in which Tao is endogenously expressed with a Venus tag (Tao-kiVenus)[34] and performed an immunostaining of Msn. Tao-kiVenus and Msn colocalized in clusters at the onset and mid-migration stages, particularly at the cluster periphery (Fig.2A,B), where Msn phosphorylates Moe. [14]. Additionally, colocalization was also observed in vesicular structures (Fig.2A, arrowheads).

**Figure 2:**
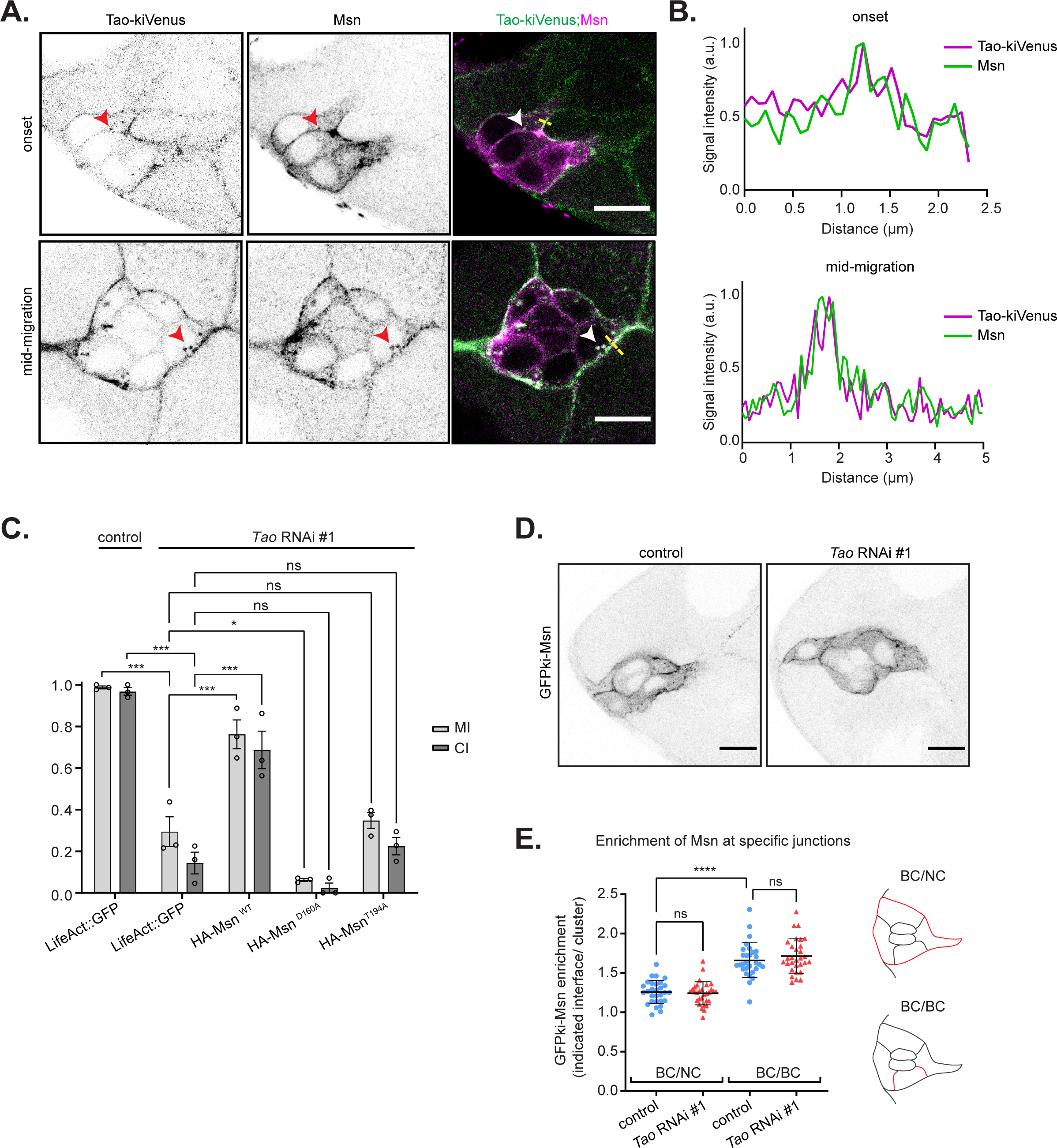
Tao activates Msn but does not regulate its localization (A) Representative confocal images showing the localization of Tao and Msn in BCs at onset and mid-migration. Their colocalization in vesicles is highlighted by red arrowheads in separated channels (shown as inverted greyscale images) and white arrowheads in merged images. Colocalization appears in white in merged images by the superposition of the Tao-kiVenus signal in green and Msn signal in magenta. Scale bar: 10µm. (B) Line scan indicates the colocalization between Tao-kiVenus (green) and Msn (magenta) at the periphery of the cluster (yellow dashed line) (C) Migration and completion indexes of BC migration determined in clusters expressing control RNAi or *Tao* RNAi together with the expression of LifeAct::GFP, a wild type form of Msn (Msn^WT^), a kinase dead form of Msn (Msn^D160A^), and a phosphodeficient form of Msn (Msn^T194A^), as indicated. (n = 145, 87, 72, 52 and 72 respectively; N=3, ns: not significant, *: p<0.05, ***: p<0.001, One-Way ANOVA, Dunnett’s correction) (D) Representative images showing the localization of GFPki-Msn at onset of migration in control and Tao-depleted clusters, displayed as inverted greyscale images. Scale bar: 10µm. (E) Quantification of the enrichment of GFPki-Msn at inner junctions (BC/BC interface) and at the periphery of the cluster (BC/NC interface) at the onset of migration in control and Tao-depleted clusters. Schemes on the right indicate in red the interfaces quantified. (n = 29 and 30 respectively; N=3, ns: not significant, ****: p<0.0001, One-Way ANOVA, Sídák’s correction). All error bars represent SEM.

Finally, we conducted rescue experiments in Tao-depleted background to verify whether the migration defects observed upon Tao depletion resulted from a lack of Msn activation (Fig.2C). We observed that the cluster migration, blocked by *Tao* RNAi expression, was significantly restored by overexpressing wild-type Msn (HA-Msn). In contrast, the expression of HA-Msn^T194A^, a phosphodeficient mutant form of Msn that cannot be phosphorylated by Tao, failed to rescue the migration delay. Similarly, the catalytically inactive mutant form of Msn (HA-Msn^D160A^) did not rescue the delay. To confirm that the differences were not due to variable expression, we verified that all three Msn constructs were expressed at similar levels (Fig.S2E,F). These data demonstrate that Tao acts as an upstream kinase of Msn to regulate BC migration.

Next, we investigated whether Tao activity towards Msn regulates its localization at the periphery of the cluster, which is crucial for Moe phosphorylation [14]. To explore this, we combined the expression of *Tao* RNAi with the GFPki-Msn allele. Depletion of Tao neither affected the enrichment of GFPki-Msn at the periphery of the cluster nor at inner junctions during the onset of migration (Fig.2D,E). Therefore, our findings suggest that while Msn activation depends on Tao, its localization at the cluster periphery is independent of Tao. Our results are consistent with previous observations in Drosophila enteroblasts, where Tao regulates the phosphorylation of Msn at T194 but does not impact its association with the membrane [21].

### Msn localization depends on its CNH domain, which interacts with Rap2l

To gain insights into the mechanism regulating the localization of Msn, we expressed different mutant forms of HA-tagged Msn in BCs using the s*lbo*-Gal4 driver. The catalytically inactive mutant form (HA-Msn^D160A^) and the phosphodeficient mutant form (HA-Msn^T194A^) of Msn localize at the cell cortex like wild-type Msn (HA-Msn^WT^). These results indicates that neither the kinase activity of Msn nor its activation by Tao are required for Msn recruitment at the cluster periphery, confirming our finding that Tao is not required to localize Msn. Interestingly, the truncated form of Msn lacking the CNH domain (HA-Msn^ΔCNH^) was exclusively found in the cytoplasm (Fig.3A), suggesting that the cortical localization of Msn depends on its CNH domain.

**Figure 3:**
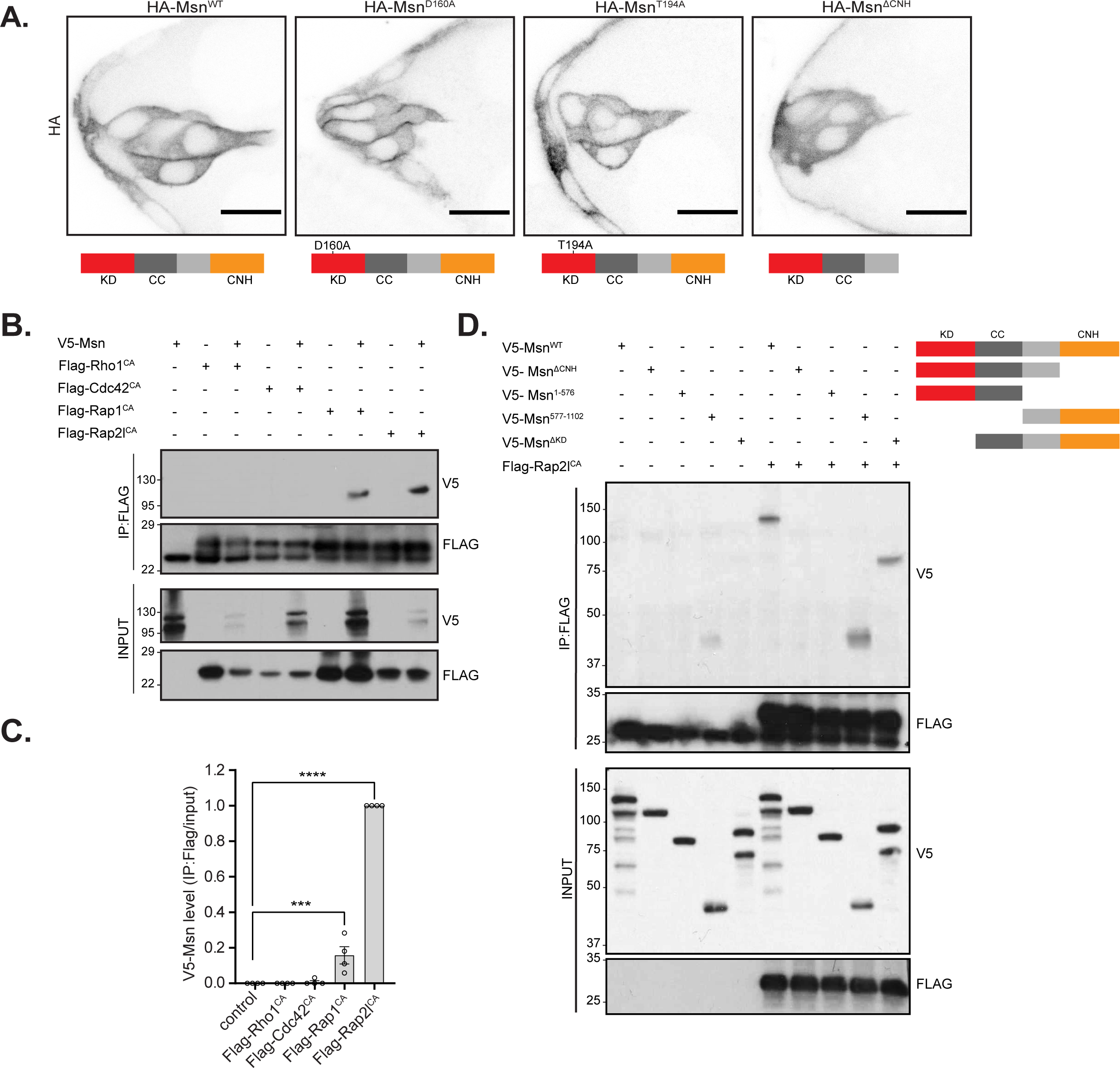
Msn localization depends on its CNH domain which interacts with Rap2l. (A) Representative confocal images showing the localization of a wild-type form of Msn (HA-Msn^WT^), a kinase dead form of Msn (HA-Msn^D160A^), a phosphodeficient form of Msn (HA-Msn^T194A^) and a form of Msn deleted for the CNH domain (HA-Msn^ΔCNH^) at onset BC clusters. The HA channel is displayed as inverted greyscale images and schemes on the bottom represent Msn protein structure with the respective mutations and truncations. KD - kinase domain, CC-coiled-coil and CNH - CNH domain. Scale bar: 10µm (B) Representative Western blot images of immunoprecipitated Flag-tagged constitutively active (CA) GTPases co-expressed with V5-tagged Msn in Drosophila S2 cells. KD - kinase domain, CC-coiled-coil and CNH - CNH domain. (C) Ratio of V5-Msn blot intensity at Flag-immunoprecipitated and input samples for each small GTPase expressed from three independent assays. (***: p<0.001, ****: p<0.0001, One-Way ANOVA, Dunnett’s correction) (D) Representative Western blot images of immunoprecipitated Flag-tagged constitutively active (CA) Rap2l^CA^ co-expressed with V5-tagged truncated forms of Msn in Drosophila S2 cells. A representation of each truncated form of Msn is displayed at the left. All error bars represent SEM.

The CNH domain is known to bind active Rho and Rap small GTPases [22, 23]. In mammals, the Msn ortholog MAP4K4 acts downstream of the small GTPase Rap2, and their interaction is mediated by the CNH domain [35]. To investigate potential upstream regulators that may impact Msn localization through binding its CNH domain, we co-expressed a V5-tagged version of Msn and a limited set of Flag-tagged constitutively active (CA) Rho and Rap small GTPases in Drosophila S2 cells and immunoprecipitated the different small GTPases with an anti-Flag antibody. We found that Msn co-immunoprecipitated with the CA forms of Rap1 and Rap2l, showing a strong preference for Rap2l (Fig.3B,C). We did not observe any interaction between Msn and the CA forms of Rho1 and Cdc42, however Msn expression levels were variable, and as such, we cannot entirely exclude a potential interaction between Msn and those small GTPases.

Next, we focused on the binding to Rap2l, as it is the strongest. To map the domain of Msn involved in this interaction, we co-expressed Flag-Rap2l^CA^ along with V5-tagged truncations of Msn. The interaction was abrogated in the two truncations lacking the CNH domain, Msn^ΔCNH^ and Msn^1-576^, indicating that Rap2l interacts with Msn through the CNH domain (Fig.3D). Overall, our findings reveal that the CNH domain is required for Msn recruitment at the cell cortex and interacts with Rap2l.

### Rap2l regulates Msn localization in BCs

After identifying Rap2l as a new interactor of Msn, we wondered if Rap2l could be a regulator of Msn during BC migration. First, we determined if Rap2l is necessary for the cluster migration. We used the *c306*-Gal4 driver to express two independent non-overlapping dsRNA against *Rap2l*. Rap2l depletion resulted in a noticeable delay in the migration indicating that Rap2l is indeed required for BC migration (Fig.4A). The two different RNAis against *Rap2l* significantly reduced Rap2l mRNA expression levels (Fig.S3A) and produced similar phenotypes (Fig.4A, D-F).

**Figure 4:**
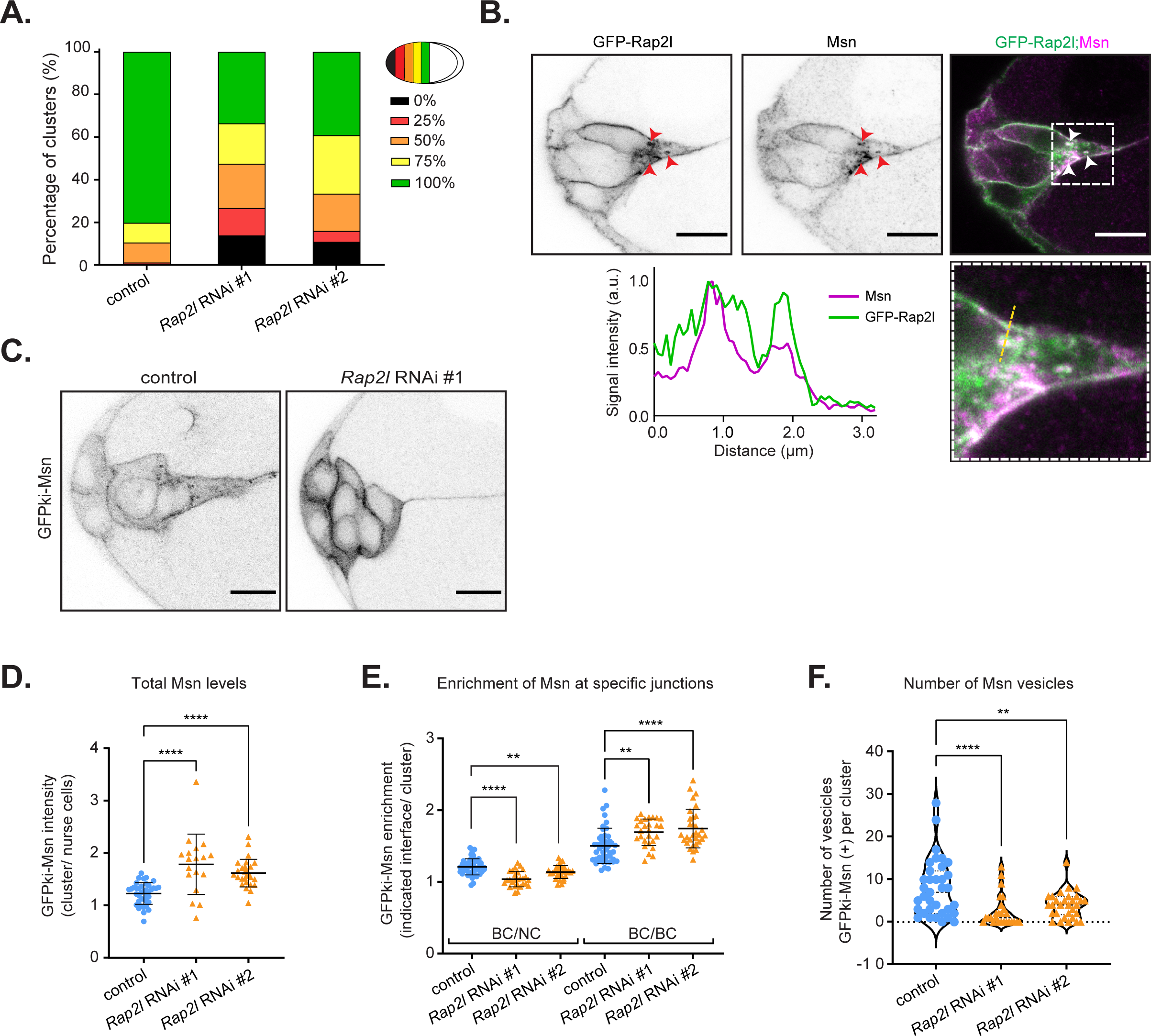
Rap2l is required for BC migration and regulates Msn distribution in BCs. (A) Quantification of the percentage of BC clusters at each sector in stage 10 egg chambers after expression of a control RNAi or two independent dsRNA against *Rap2l*. (n= 86, 116 and 138, respective to the histogram order. N=3) (B) Representative confocal images of the localization of GFP-Rap2l and Msn at the onset of migration. Their colocalization in vesicles is highlighted by red arrowheads in separated channels (shown as inverted greyscale images) and white arrowheads in merged images. Colocalization appears in white in merged images by the superposition of the GFP-Rap2l signal in green and Msn signal in magenta. At the bottom, high magnification image and line scan demonstrate similar distribution of mean intensity of Rap2l and Msn at vesicles and the periphery of the cluster (yellow dashed line). Scale bar: 10µm (C) Representative confocal images showing the localization of GFPki-Msn at onset of migration in control and Rap2l-depleted clusters, as inverted greyscale images. Scale bar: 10µm. (D) Quantification of the ratio of GFPki-Msn mean intensity in the entire cluster, normalized to the signal between nurse cells, in control and Rap2l-depleted clusters (n = 39, 18 and 27 respectively; N=3, ****: p<0.0001, One-Way ANOVA, Dunnett’s correction) (E) Quantification of the enrichment of GFPki-Msn at inner junctions (BC/BC interface) and at the periphery of the cluster (BC/NC interface) at the onset of migration in control and Rap2l-depleted clusters (n = 45, 27 and 31 respectively; N=3, ****: p<0.0001, One-Way ANOVA, Dunnett’s correction). (F) Quantification of the number of GFPki-Msn positive vesicles per cluster at the onset of migration in control and Rap2l-depleted clusters. (n = 35, 26 and 30 respectively; N=3, **: p<0.01, ****: p<0.0001, One-Way ANOVA, Dunnett’s correction). All error bars represent SEM.

To gain more insights into the function of Rap2l, we examined its subcellular localization by overexpressing a GFP-fused version of Rap2l (GFP-Rap2l). We found a predominant cortical localization of GFP-Rap2l at the periphery of the cluster and inner junctions. Furthermore, we detected GFP-Rap2l in punctate structures, suggesting a vesicular localization (Fig.4B). These observations corroborate with previous studies, in which active Rap2 traffics from the plasma membrane to endosomes [36, 37].

Co-staining with the Msn antibody demonstrated colocalization between Msn and GFP-Rap2l at the cell cortex as well as in vesicular structures (Fig.4B). A similar observation was made when we combined expression of HA-tagged Rap2l (HA-Rap2l) with GFPki-Msn (Fig.S3B). Together with the biochemistry evidence of Rap2l and Msn interaction, this suggests that Rap2l and Msn interact in those specific subcellular compartments.

Our findings show that Rap2l interacts with the CNH domain of Msn, which is required to recruit Msn to the cortex. Based on this, we hypothesized that Rap2l might regulate the localization of Msn. To investigate this, we knockdown *Rap2l* in GFPki-Msn flies. Surprisingly, we observed that Msn remained enriched at the cell cortex even after depleting Rap2l (Fig.4C), suggesting that Rap2l is not required to recruit Msn at the cell cortex.

However, we did observe significant differences in the localization of Msn compared to control conditions. These changes were specific to Msn, as Tao-kiVenus was unaffected by Rap2l depletion (Fig.S3C-E). Firstly, there was a significant increase in the overall levels of GFPki-Msn in clusters expressing *Rap2l* RNAi (Fig.4D). *Msn* mRNA levels in control and *Rap2l* RNAi-expressing flies were comparable (Fig.S3F) suggesting that the elevated levels of Msn protein were not due to increased transcription. Overall, these findings suggest that Rap2l may regulate Msn stability on BCs.

Secondly, we observed that following Rap2l depletion, Msn was distinctly distributed, with reduced enrichment at the periphery of the cluster and increased levels at the inner junctions (Fig.4E). Additionally, there was an apparent reduction in Msn vesicles after Rap2l depletion (Fig.4F), indicating that Rap2l may be required to transport Msn in a vesicular compartment. In contrast, when Rap2l was overexpressed, although there was no detectable change in vesicle number and total levels of Msn (Fig.S3G-I), the distribution of GFPki-Msn was opposite to what observed after Rap2l depletion. Rap2l overexpression resulted in increase on Msn enrichment at the periphery of the cluster and a slight, non significant, decrease on Msn levels at the inner junctions (Fig.S3G,J).

Considering previous work indicating that the distribution of Msn at the periphery of the cluster relies on vesicular trafficking [14], our findings suggest that Rap2l may regulate the distribution of Msn within the cluster by modulating its trafficking.

### Rap2l regulates the trafficking of Msn into late endosomes

We previously described that Msn was present in recycling endosomes and its localization at the periphery of the cluster depended on Rab11, which regulates the recycling pathway [14]. However, the localization of Msn in others endocytic compartments has not been investigated. To extend our previous analysis, we co-stained Msn with markers of diverse endosomal compartments in control clusters.

First, we observed little colocalization of Msn with early endosomal markers (GFP-Rab5) [38] (Fig.S4A,B). Proteins within the early endosomes can be sorted toward two distinct pathways: recycling or degradation. In the recycling pathway, the proteins are transported back to the plasma membrane for reuse. In contrast, the degradative pathway involves the transport of the protein to late endosomes, which can fuse with lysosomes where degradation occurs [39].

Using a YFP::Msn, a fluorescent protein trap in the *msn* gene, we previously detected the presence of Msn in recycling endosomes [14], using Sec15-mCherry [40] or Rab11-mCherry as markers. Yet, using the antibody against Msn that we validated in this work (Fig,S2C,D), we could not observe a significant colocalization with the marker associated with recycling endosome GFP-Sec15 (Fig.S4C). However, we observed an evident accumulation of Msn inside late endosomes labelled with GFP-Rab7 (Fig.5A,B). These findings indicate that the observed vesicular pool of Msn localizes mainly in late endosomes in BCs.

**Figure 5:**
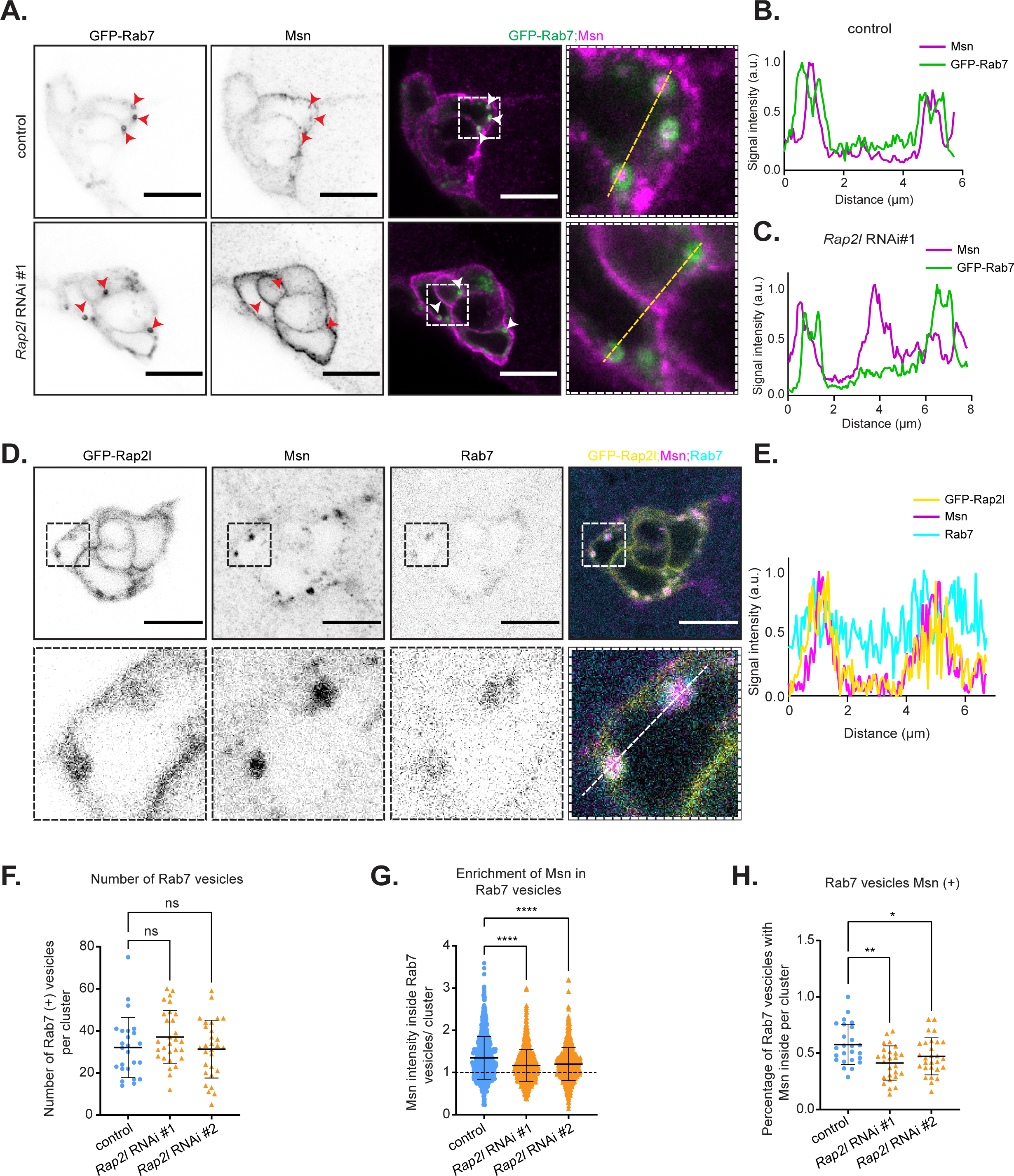
Rap2l regulates the trafficking of Msn into late endosomes. (A-C) Representative confocal images of co-labelling of Msn by immunostaining and late endosomes (marked with GFP-Rab7) in control and Rap2l-depleted clusters. The localization in vesicles is highlighted by red arrowheads in separated channels (shown as inverted greyscale images) and by white arrowheads in merged images. A high magnification image is presented on the right. Scale bar: 10μm. Quantification of the fluorescence signal of Msn and Rab7 by “line scan” (along the yellow, dashed line shown in A) in control (B) and Rap2l-depleted cluster (C). (D) Representative confocal images showing the localization of GFP-Rap2l, Msn and Rab7 in BCs. Their colocalization in vesicles is highlighted in high magnification images of separated channels (shown as inverted greyscale images) and in merged images by superposition of the GFP-Rap2l signal in yellow, Msn signal in magenta and Rab7 signal in cyan. Scale bar: 10µm (E) Quantification of the fluorescence signal of Rap2l, Msn and Rab7 by “line scan” (along the yellow, dashed line shown in D). (F) Number of Rab7 positive vesicles in control clusters and cluster depleted for Rap2l. (n = 25, 29 and 29 respectively; N=3, ns= not significant, One-Way ANOVA, Dunnett’s correction) (G) Intensity of Msn signal within Rab7-positive structures normalized by the overall intensity of Msn in control clusters and clusters depleted of Rap2l (n = 803, 994 and 899 respectively; N=3, **** p<0.0001, One-Way ANOVA, Dunnett’s correction) (H) Percentage of Rab7 vesicles with Msn enriched (enrichment > 1.2) per cluster in control and Rap2l-depleted cluster (n = 25, 26 and 28 respectively; N=3, * p<0.05, ** p<0.01, One-Way ANOVA, Dunnett’s correction). All error bars represent SEM.

Due to the interaction between Rap2l and Msn in Drosophila cells and the presence of both proteins in vesicular compartments, we questioned if Rap2l is involved in the trafficking of Msn. Previous studies have reported the localization of Rap2l orthologs in endosomes and their function in sorting proteins, including MINK and TNIK, two homologs of MAP4K4 [24, 37, 41, 42]. Because Msn is mainly found in late endosomes, we first investigated whether Rap2l localizes at late endosomes in BCs. Co-staining of BCs expressing GFP-Rap2l with Rab7 and Msn antibodies shows that Rap2l and Msn colocalized in late endosomes (Fig.5D,E).

Since Rap2l depletion led to a reduction of GFPki-Msn positive vesicles (Fig.4F), we next examined if Rap2l promotes the localization of Msn in late endosomes. We stained Msn in clusters expressing *Rap2l*

RNAis and GFP-Rab7 (Fig.5A,C). While we did not observe significant changes in number of Rab7 vesicles after Rap2l depletion (Fig.5F), we observed a decrease in the levels of Msn detected inside Rab7 vesicles. Specifically, there was a decrease in the percentage of Rab7 vesicles containing Msn in clusters expressing *Rap2l* RNAi (Fig.5A,G-H). These findings indicate that Rap2l promotes the localization of Msn in late endosomes without affecting the late endosome compartment per se.

Overall, our observations suggest that Rap2l promotes the targeting of Msn into late endosomes for degradation. In the absence of Rap2l, Msn is not properly degraded in the endolysosomal pathway. Instead, it accumulates in the cytoplasm and at inner junctions (Fig.6). Finally, our findings suggest that the balance between Msn at inner junction and at the periphery of the cluster is regulated by endolysosomal degradation.

**Figure 6:**
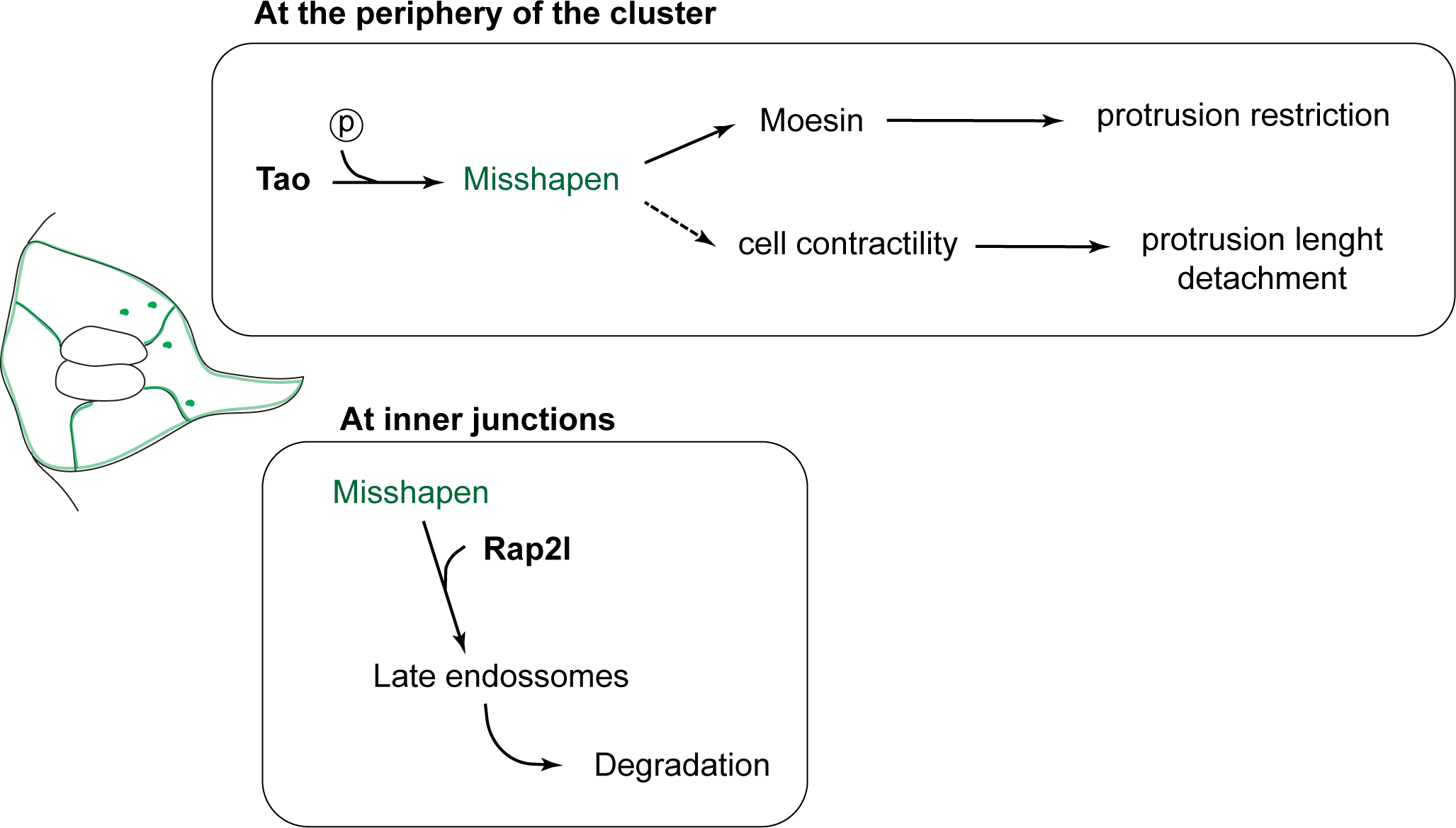
Proposed model representing how Tao regulates BC migration by activating of Msn at the periphery of the cluster while Rap2l promotes Msn degradation at inner junctions.

## DISCUSSION

During collective cell migration, coordination of protrusion formation and contractility is essential to promote directional and efficient movement. Previous studies by us and others have characterized a supracellular actomyosin structure situated at the periphery of the cluster which plays a major role in this coordination [13-15, 17]. In particular, we found that Msn is fundamental for the formation and functionality of this structure, by phosphorylating Moe and promoting contractility [14]. In the present study, we demonstrate that a dual regulation, involving Tao kinase and Rap2l, promotes the activity of Msn and favours its subcellular localization to the periphery of the cluster.

In our search for a potential activator of Msn, we investigated the role of Tao in BC migration. We found that Tao is required for BCs, as its depletion strongly impairs the cluster migration. Furthermore, the loss of Tao results in a phenotype similar to the loss-of-function phenotype observed in Msn, including defects in cluster detachment and protrusion restriction. These findings align with previous work in Drosophila midgut, which identified Tao as an upstream kinase of Msn [21]. Moreover, the rescue of the Tao depletion phenotype through the overexpression of Msn^WT^ reinforces the notion that Msn is downstream of Tao in BCs. This cascade involving Tao and Msn is likely to be conserved in mammals as MAP4K4, the ortholog of Msn, was shown to be required for the phosphorylation of the transcription factor YAP mediated by the Tao kinase ortholog TAOK-1 [43].

Our work does not exclude that Tao has additional functions in BCs beyond phosphorylating Msn. In previous studies, Tao was shown to act as a negative regulator of microtubules stability and an upstream kinase of Hippo [26, 28, 44]. To our knowledge, the role of microtubules in BCs has not been extensively studied. Based on the literature [45], the phenotype we observe in BC clusters depleted of Tao appears to be more significant than what would be anticipated for microtubule stabilization. Regarding the Hippo pathway, it has been demonstrated that Hippo inhibits the actin regulator Enabled in BCs, leading to impaired actin polymerization at BC contacts [46]. In our experiments, we did not observe any apparent differences in actin distribution following Tao depletion suggesting that Tao is not a major regulator of the Hippo pathway. However, it would be valuable to formally confirm that Tao has any impact on Hippo in our particular model.

In BCs, the actomyosin supracellular structure surrounding the cluster blocks the formation of protrusion in follower cells [16, 17]. This requires the phosphorylation of Moe, therefore Msn is expected to be active at the periphery of the cluster. Accordingly, Tao was found at the periphery of the cluster and hence may activate Msn at this location. Initially, we tested if Tao would also recruit Msn to the periphery; however, similar to findings in the midgut [21], our study shows that Tao phosphorylation does not regulate the localization of Msn. Therefore, we investigated whether Msn localizes at the periphery through another mechanism, considering possible interactions with other membrane proteins. Trying to determine which domain of Msn could be related to its localization, we found that its CNH domain is essential for its recruitment to the cortex. Again, this appears to be conserved as we recently found that MAP4K4 also relies on its CNH domain for localization at cell-cell junctions [25].

Looking for regulators of Msn localization, we discovered that Rap2l interacts with Msn through its CNH domain. In Drosophila, limited information was available regarding Rap2l and its function was still unknown [47, 48]. However, the ortholog of Rap2l, Rap2 has been shown to bind to orthologs of Msn, MAP4K4, MINK and TNIK, through their CNH domain [24, 49]. Unexpectedly, despite the interaction between Rap2l and Msn, depleting Rap2l did not impair the localization of Msn at the membrane. These results indicate that Rap2l may not be directly responsible for the recruitment of Msn at the membrane. Nevertheless, it is still possible that other small GTPases, including Rap1, which binds to Msn with lower affinity compared to Rap2l, could recruit Msn at the membrane, compensating for the absence of Rap2l. Further testing and validation are necessary to test this hypothesis.

Although Msn localization to the membrane was not abrogated by Rap2l depletion, we did observe significant changes in Msn distribution within the cluster. Specifically, there was an overall increase of Msn levels in the entire cluster, particularly at the inner junctions, along with a pronounced decrease of Msn within vesicles. Specifically, we observed reduced Msn levels in late endosomes following Rap2l depletion. In mammals, Rap2 regulates the localization of MINK and TNIK to endosomes in neurons [24], however, the precise mechanism by which Rap2 regulates the presence of Msn orthologs in endosomes remains unclear. We speculate that Rap2l serves as an accessory protein that facilitates the internalization of Msn in the endolysosomal pathway to promote its degradation.

Finally, this raises questions regarding the necessity of Msn degradation in BC clusters. By monitoring the distribution of Msn at different stages of migration, we noted significant changes in the distribution of Msn between onset and mid-migration clusters. Specifically, we found that Msn is slightly more enriched at the inner junctions at the onset stage but is preferentially distributed at the periphery of the cluster at mid-migration. BCs retain certain levels of cell polarity inherited from the follicular cells from which they originate. As the cluster transits from the onset stage to mid-migration, it undergoes a not fully understood 90° rotation, resulting in changes in the distribution of specific polarity markers. For instance, actin, which is initially localized in the apical part of onset clusters, becomes more enriched at the periphery of migrating clusters [50]. In follicular cells, Msn is enriched at both the apical and lateral membranes (Fig.S2C, arrowheads). We speculate that during the initial stages of migration, the cluster likely retains the distribution of Msn inherited from follicular cells, with Msn localized to the inner junctions. However, as migration progresses, this “follicular cell distribution” appears to be lost and Msn becomes predominantly enriched at the peripheral membrane. Our results indicates that Rap2l is critical in this process, possibly by targeting the pool of Msn at inner junctions for endolysosomal degradation (Fig.6).

Overall, our work reveals two distinct regulatory mechanism acting on different domains of Msn, ensuring Msn proper distribution and activation levels. It would be interesting to investigate if these regulatory mechanisms are coordinated and interconnected. Furthermore, since the regulatory mechanisms identified in our study are likely conserved in mammals, it would be interesting to explore if TAOK and Rap2 have similar functions in the context of MAP4K4, which is often found overexpressed in solid tumors and associated to pro-metastatic behaviors [51–54].

## MATERIAL AND METHODS

### Drosophila genetics

Fly stocks were maintained at 25°C with 70% relative humidity. Crosses were performed at the same temperature, and flies were subsequently incubated at 29°C for 48 hours before dissection. In the case of the protrusion analysis of Tao knockdown, female flies were kept at 25°C prior to dissection, as the protrusion phenotype was more pronounced at lower temperatures. Knockdown and rescue experiments utilized the *c306*-Gal4 driver, whereas overexpression assays were conducted using the *slbo*-Gal4 driver. Both drivers are specifically expressed in BCs.

The three *Tao* RNAi lines used were UAS-*Tao* RNAi (RNAi #1, #17432), UAS-*Tao* RNAi (RNAi#2, #107645) obtained from the Vienna Drosophila RNAi Center, along with UAS-*Tao* RNAi (RNAi #3, # 35147) acquired from the Bloomington Stock Collection. For the *Rap2*l RNAi lines, we used UAS-*Rap2l* RNAi (RNAi #1, #45228) and UAS-*Rap2l* RNAi (RNAi #2, #107745), both obtained from the Vienna Drosophila RNAi Center. As a control, we used the UAS-*mCherry* RNAi line (#35785) from the Bloomington Stock Collection. Other stocks from the Bloomington Stock Collection included *c306*-GAL4 (#3743), *UAS-Lifeact::GFP* (#35544), UAS-*msn* RNAi (#28791) and UAS-lacZ (#8529).

Tao^eta^ FRT19A/FM6 was obtained from the group of Veit Riechmann (Universität Heidelberg) and Tao-kiVenus was provided by the group of Kieran Harvey (University of Melbourne). GFPki-Msn and UAS-HA-Msn^T194A^/TM2a were obtained from the group of Tony IP (University of Massachusetts Medical School). UAS-GFP-Rab5/TM3 and UAS-GFP Rab7/CyO were acquired from the group of M. Gonzalez-Gaitan (Université de Genève) and UAS-eGFP-sec15/TM3 and Hs-Gal4/Cyo were donated by the group of J.A. Knoblich (Institute of Molecular Biotechnology of the Austrian Academy of Sciences).

The following lines: UAS-HA-Msn^WT^, UAS-HA-Msn^D160A^, UAS-HA-Msn^ΔCNH^ [14], FRT19A, hsFLP, TubGal80 /FM7; *Slbo*-GFP /CyO and UAS-Rab11-mCherry*/*CyO [15] were previously generated in our laboratory. Genome Prolab generated the transgenic flies UAS-GFP-Rap2l and UAS-HA-Rap2l by injecting a pUAS-attB plasmid containing the specific tag and Rap2l sequence (cloning procedure explained below).

To generate genetic mosaics of Tao^eta^, we performed crosses between Tao^eta^ FRT19A and FRT19A, hsFLP, TubGal80 /FM7; *Slbo*-GFP /CyO flies. Heat shock was used to induce BC clones by immersing the tube with pupae in 37°C water bath for one hour for three consecutive days before hatching and one additional day after hatching. Following the heat shock treatment, the female flies were incubated for two days at 25°C with yeast before dissection.

### Cloning

Rho1 and Cdc42 sequences were subcloned from pOT2-Rho1 (clone GH20776) and pOT2-Cdc42 (clone HL08128) plasmids, respectively (gift from Vincent Archambault, IRIC, Montreal). Rap2l sequence was obtained from a pFlc-1-Rap2l (clone RE63021) plasmid (gift from Vincent Archambault, IRIC, Montreal). Rap1 sequence was cloned from a pMet-3xHA-Rap1 (gift from Marc Therrien, IRIC, Montreal). The following primers were used to amplify the respective sequences and insert them into a Gateway vector pDONR221 generating pEntry-Rho1, pEntry Cdc42, pEntry-Rap2l and pEntry-Rap1.

For Rho1: 5’ GGGGACAAGTTTGTACAAAAAAGCAGGCTTCATGACGACGATTCGCAAGAAATTG 3’ and 5’ GGGGACCACTTTGTACAAGAAAGCTGGGTCTTAGAGCAAAAGGCATCTGGTCTT 3’ For Cdc42: 5’ GGGGACAAGTTTGTACAAAAAAGCAGGCTTCATGCAAACCATCAAGTGGT 3’ and 5’ GGGGACCACTTTGTACAAGAAAGCTGGGTCTTATAAGAATTTGCACTTCCTTTTCTTTGT 3’ For Rap2l: 5’ GGGGACAAGTTTGTACAAAAAAGCAGGCTTCATGCGCGAATTCAAAGTTGTTGTG 3’ and 5’ GGGGACCACTTTGTACAAGAAAGCTGGGTCCTATAAAAGCGTACAACAACAGTA 3’ For Rap1: 5’ GGGGACAAGTTTGTACAAAAAAGCAGGCTTCATGCGTGAGTACAAAATCGTGGTC 3’ and 5’ GGGGACCACTTTGTACAAGAAAGCTGGGTCTTATAGCAGAACACATAGGGACTT 3’ To generate the constitutively active forms of each GTPase, we performed mutagenesis using the following primers: For Rho1^CA^ (G14V): 5’ GTCGGCGACGTCGCCTGC 3’ and 5’ AATTACCAATTTCTTGCGAATCGTCGTCATG 3’ For Cdc42^CA^ (G12V): 5’ TCGGCGACGTCGCCGTGG 3’ and 5’ CGACCACGCACTTGATGGTTTGCAT 3’ For Rap2l^CA^ (G12V): 5’ CGGGTCGGTCGGAGTTGGTAAA 3’ and 5’ AGCACAACAACTTTGAATTCGCGCAT 3’ For Rap1^CA^ (G12V): 5’ TTGGAAGCGTCGGCGTGG 3’ and 5’ GGACCACGATTTTGTACTCACGCAT 3’ To obtain pA-Flag-Rho1^CA^, pA-Flag-Cdc42^CA^, pA-Flag-Rap2l^CA^, pA-Flag-Rap1^CA^ vectors, the respective sequences were inserted into the pAc5 gateway cassette N-ter 3*flag-tag (pAFW, RRID:DGRC_1111). These vectors were subsequently used for the co-immunoprecipitation assay. Plasmids pMT-V5-Msn, pMT-V5-Msn^ΔCNH^, pMT-V5-Msn^1-576^, pMT-V5-Msn^576-1102^ and pMT-V5-Msn^ΔKD^ were provided by Marc Therrien, IRIC, Montreal.

For the expression of Rap2l in flies, Rap2l was subcloned from our pEntry-Rap2l plasmid into the pUAS-AttB plasmid (from Marc Therrien, IRIC, Montreal). For pUAS-HA-Rap2l, the HA sequence was added to the Rap2l sequence separated by AatII and AvrII restriction site using the primer: 5’ATGTACCCATACGATGTTCCAGATTACGCTGACGTCCGCGAATTCAAAGTTGTTG 3’. Then, HA-Rap2l was cloned to pUAS-AttB plasmid using the Gibson technique. The primers used for this step were: 5’ GCGCGGCCGCCATGTACCCATACGATGTTCCAGATTACGCTCGCGAATTCAAAGTTGTTG 3’ and 5’ GCGGGGCGCGTGGTACCCTATAAAAGCGTACAACAACAGTA 3’. For pUAS-GFP-Rap2l, The GFP sequence was subsequently cloned into the pUAS-HA-Rap2l plasmid replacing the HA tag. This was done using the AatII and AvrII restriction sites. All constructs were verified by Sanger sequencing.

### mRNA and protein extraction

For mRNA and protein extraction, crosses were made using the Hs-Gal4 as the driver. Females were heat-shocked at 37 °C for one hour and then kept at 29 °C for 72 hours before dissection. mRNA was extracted by using the RNeasy Mini kit (Quiagen) in female flies without ovaries. Protein extraction was performed from the whole fly using the lysis buffer (50 mM Tris-HCl pH 7.5, 150 mM NaCl, 1% Triton X-100, 10% glycerol, 1 mM EDTA) as instructed in [55].

### Co-immunoprecipitation

For the co-immunoprecipitation assay, Drosophila S2 cells were cultured in Excel media without fetal bovine serum (FBS) and co-transfected with Effectene (Qiagen). The expression of the copper-inducible transgenes was induced by adding CuSO4 (700 µM) for 36 hours. After that, the cells were lysed on ice for 20 minutes using lysis buffer (50 mM Tris-HCl pH 7.5, 150 mM NaCl, 1% Triton X-100, 10% glycerol, 1 mM EDTA), supplemented with 20 μM leupeptin, aprotinin (0.15 U/mL), 1 mM PMSF, a phosphatase inhibitor cocktail (Sigma-Aldrich), and 1 mM sodium orthovanadate. The cell lysates were centrifuged at 14,000 rpm at 4°C for 10 minutes, and the resulting supernatant was collected. Protein concentrations were determined using the BCA Protein Assay Kit (Thermo Fisher Scientific).

To immunoprecipitated the pA-Flag tagged constitutively active small GTPases, the cell lysates were pre-incubated with 1 µL of anti-Flag M2 antibody (Sigma-Aldrich) for 2 hours at 4°C. Subsequently, 10 μL of protein A/G beads were added, and the mixture was incubated overnight. The beads were then washed at least three times with cold lysis buffer and boiled in 2X sample loading buffer (100 mM Tris-HCl pH 6.8, 4% SDS, 0.2% bromophenol blue, 20% glycerol, 200 mM β-mercaptoethanol) for 5 minutes before performing SDS-PAGE.

For immunoblotting analysis, lysates or immunoprecipitated proteins were separated on SDS-PAGE gels and transferred onto PVDF membranes (Bio-Rad). The membranes were blocked with skim milk (5%) or 2% BSA (Sigma-Aldrich) for 1 hour and incubated with the corresponding primary antibodies for 2 hours at room temperature (RT) or overnight at 4°C. The primary antibodies used were as follows: anti-FLAG M2 from mouse (1:5000; Sigma-Aldrich; #F1804), anti-V5 Tag from mouse (1:5000; Invitrogen; #R960-25), anti-HA from rabbit (1:1000; Invitrogen; #71-5500), and mouse monoclonal anti-Actin at 1:10000 (Millipore (C4), MAB1501).

For the secondary antibodies, we used anti-mouse-HRP (1:5000; Jackson ImmunoResearch Laboratories Inc.; #115-035-062), anti-mouse-HRP light chain (1:5000; Jackson ImmunoResearch Laboratories Inc.; #115-035-174), and anti-rabbit-HRP light chain (1:5000; Jackson ImmunoResearch Laboratories Inc.; #211-032-171). They were incubated at RT for 1 hour, and the membranes were visualized using an X-ray film in a dark room. Protein levels were quantified from Western Blot film image using Analyze>Gel tool of Fiji (Image J).

### Immunofluorescence

Ovaries were dissected from young adult females in PBS 1X and fixed by incubation in 4% paraformaldehyde diluted in PBS for 20 minutes. Afterwards, the ovaries were washed 3 times with 200 µL of wash solution (Triton X-100, 0.3% in PBS) and incubated in 200µL of block solution (2% BSA + Triton X-100 0.3% in PBS) for 2 hours at RT with agitation. Next, the ovaries were incubated overnight at 4°C with the respective primary antibody diluted in block solution. The following antibodies were used at the indicated dilutions: rabbit polyclonal anti-phospho-Ezrin (Thr567)/Radixin (Thr564)/Moesin (Thr558) at 1:50 (Cell Signaling Technology, #3141), rabbit anti-Msn M1 at 1:750 (gift from Thomas Moss, University of Laval), mouse anti-HA at 1:100 (gift from Marc Therrien, IRIC, University of Montreal) and mouse anti-rab7 at 1:15 (DSHB, AB_2722471).

After the primary antibody incubation, the ovaries were washed twice briefly and twice 20 minutes under agitation in the wash solution. They were then incubated with the secondary antibody for 2 hours at RT. The secondary antibodies used were anti-rabbit conjugated to Alexa Fluor 555 at 1:250 (Cell Signaling Technology #4413), anti-mouse conjugated to Alexa Fluor 555 (Cell Signaling Technology #4499) at 1:250 and anti-mouse conjugated to Alexa Fluor 635 (Invitrogen #A31575) at 1:100. Following the secondary antibody incubation, the ovaries were washed again twice briefly and twice for 20 minutes. Finally, they were mounted on slides using Vectashield (H-1000, Vector Laboratories). For additional staining, Alexa Fluor 555-labeled Phalloidin at 1:250 (Invitrogen, #A34055) and Alexa Fluor 647-labeled Phalloidin (Invitrogen, #A22287) at 1:100 were used to visualize F-actin, and DAPI (D8417-10MG, Sigma) was used at 1:10,000 to stain DNA. These additional stains were incubated together with the secondary antibody.

### Image acquisition and quantitative analysis

Images were captured using a laser scanning confocal microscope LSM 880 (Carl Zeiss) or a Leica TCS SP8 (Leica Microsystems).

The analysis and quantification of the number, angle, and length of protrusions were performed on fixed samples of LifeAct::GFP expressing clusters using Fiji (Image J). To quantify the number of protrusions, a circle was drawn around the main body of the cluster. Any actin extension beyond 4µm outside of the circle was considered a protrusion, while extensions longer than 9 µm were defined as main protrusions. For distribution analysis, the angle between a line drawn from the cell nucleus to the tip of the protrusion and the direction of migration was measured for each protruding cells. To represent the data, each protrusion was aligned on a radar map divided into eight sectors of 45 degrees each, with the leading-edge set at zero degrees and the trailing edge at 180 degrees. The lengths of protrusion were determined by measuring the distance between the nucleus and the tip of the protrusion for each protruding cells.

Quantification of pMoe and Moe fluorescence intensities was performed on images of fixed tissues as described in [56]. The mean intensity of pMoe and Moe at the periphery of the cluster was measured using three z-stack images. The LifeActGFP signal was used to define the periphery of the cluster. To account for variation in staining across egg chambers, the signal of BCs was normalized by the intensity measured on nurse cell membranes.

For the analysis of GFPki-Msn localization, the following steps were performed: mean GFPki-Msn fluorescence intensities were quantified at the peripheral membrane and at a well-defined BC-BC interface in three z-stacks images. The phalloidin staining was used to delineate the periphery of the cluster and the BC-BC interface. The enrichment of Msn was calculated by dividing the mean intensity at each location by the mean intensity of GFPki-Msn measured in the entire cluster. To account for variations in signal across egg chambers and compare the overall levels of GFPki-Msn in the cluster, the signal measure in the entire cluster was normalized by the intensity measured on the nurse cell membrane. The number of GFPki-Msn vesicles (GFPki-Msn positive puncta) was quantified by manually counting the vesicles observed in a complete z-scan of the cluster. The same methodology was used to quantify Tao-kiVenus localization.

For all these analyses, the background signal, determined outside the egg chamber, was consistently subtracted from the measured fluorescence intensities. Only images with signals within the linear range were included for quantification.

The analysis of GFP-Rab7 vesicles was conducted using the Spot plugin on Imaris (Bitplane) on a z-scan recorded with frames separated by 1µm. The intensity of GFPki-Msn at vesicles was normalized by the mean overall levels of GFPki-Msn obtained from five z-stack of each cluster. This normalization was performed to calculate the enrichment of Msn at the position of the Rab7 vesicles. To determine the percentage of Rab7 vesicles with Msn enriched inside, we quantified the number of Rab7 vesicles with Msn enrichment greater than 1.2 and divided by the total number of Rab7 vesicles in the cluster.

### Migration Quantification

BC migration was quantified from stage 10 egg chamber using an inverted microscope AxioImager (Carl Zeiss). The migration index (M.I) was calculated as a measure of the relative distance reached by the BCs compared to the end of the migration path. The M.I is determined using the following formula:

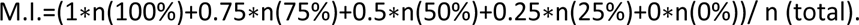

In this formula, n(100%) represents the number of egg chambers where the cluster reached the oocyte, n(75%) represents the number of egg chambers where the cluster migrated to 75% of the final distance, and so on. N (total) represents the total number of egg chambers analyzed. The Completion Index (C.I.) is calculated as the number of egg chambers where the migration was completed divided by the total number of egg chambers counted:

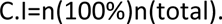

### Statistical analysis

All graphs and statistical tests were performed using GraphPad Prism software (GraphPad Software). For each analysis, we conducted a minimum of three independent experiments. Statistical analyses were carried out using unpaired Student’s t-test, non-parametric Mann-Whitney test, or One-way ANOVA test with Dunnet’s or Sikak’s correction for multiple comparisons of unpaired data. Values are presented as the mean ± standard error of the mean (SEM), and individual data points are shown on the graphs. N represents the number of independent egg chambers and N represents the number of independent experiments. P-values are indicated on the figures as follows: (*) for p<0.05, (∗∗) for p<0.01, (∗∗∗) for p<0.001 and (∗∗∗∗) for p<0.0001.

## ACKNOWLEDGMENTS

We thank the Bloomington Stock Collection and the Vienna Drosophila RNAi Center for fly stocks. We thank Tony Ip, Veit Riechmamm, Kieran Harvey, Thomas Moss, Vincent Archambault, Marc Therrien, Marcos Gonzalez-Gaitan and Jürgen Knoblich for their generosity in sharing reagents. We thank Christian Charbonneau, Malha Shami and Eloise Duramé (IRIC, Montreal) for technical assistance and the entire Emery lab for the helpful discussions and comments on the manuscript. We thank Jocelyn McDonald (Kansas State University) for helpful discussions and for sharing unpublished results about Rap GTPases. This work was supported by grants from the Canadian Institutes for Health Research to G.E. (PJT-148560 and PJT-186133), G.M.R held a doctoral scholarship from Fonds de Recherche du Quebec – Sante (FRQS).

## FIGURE LEGENDS

**Figure S1:** Expression of dsRNAs against *Tao* reduces Tao protein levels and induce defects in protrusion restriction and detachment (A) Representative images showing the intensity and distribution of Tao-kiVenus at the onset of migration in control and Tao-depleted clusters. The Tao-kiVenus channel is displayed individually as inverted greyscale images. Scale bar: 10µm (B) Quantification of the ratio of Tao-kiVenus mean intensity at the cluster periphery, normalized to the signal between nurse cells in control and Tao-depleted clusters (n= 20, 18, 11 and 12, respectively, N=3, ***: p<0.001, ****: p<0.0001, One-Way ANOVA, Dunnett’s correction) (C) Quantification of the percentage of clusters that have not detached from the follicular epithelium by stage 10 (fixed samples) (n>30; N=3; **: p<0.01, ***: p<0.001 One-Way ANOVA, Dunnett’s correction). (D) Quantification of the number of protrusions per cluster in control and Tao-depleted clusters. Error bars are SEM (n = 51, n = 43 and n=43, respectively. N=3, ****: p<0.0001, One-Way ANOVA, Dunnett’s correction). Any protrusions longer than 4 μm were manually counted using fixed samples expressing LifeAct::GFP((E) Quantification of the length of protrusions in control and Tao-depleted clusters. (n = 51, n = 43 and n=43, respectively. N=3, ****: p<0.0001, One-Way ANOVA, Dunnett’s correction) (F) Quantification of the angle between the tip of protrusions longer than 9μm and the direction of migration in control and Tao-depleted clusters. The variance between experiments is shown in the graph below (n = 28, 71 and 61, respectively; N=3 **: p<0.01, ***: p<0.001 One-Way ANOVA, Dunnett’s correction). (G) Quantification of the ratio of pMoe mean intensity at the cluster periphery, normalized to the signal between nurse cells, in cluster expressing control RNAi and *Tao* RNAi. (n= 35 and 37 and 42, respectively, N=3, ****: p<0.0001, One-Way ANOVA, Dunnett’s correction) (H) Quantification of the ratio of total Moe mean intensity at the cluster periphery, normalized to the signal between nurse cells, after expression of control RNAi and *Tao* RNAi. (n= 29, 32 and 32, respectively; N=3, ns: not significant, One-Way ANOVA, Dunnett’s correction). All error bars represent SEM.

**Figure S2:** Msn distribution changes during the migration of BC clusters. (A) Representative confocal images of the localization of GFPki-Msn at onset and mid-migration displayed as inverted greyscale images. Scale bar: 10µm (B) Quantification of the enrichment of GFPki-Msn at inner junctions (BC/BC interface) and at the periphery of the cluster (BC/NC interface) at the onset of migration and in mid-migration clusters. Schemes bellow indicate the regions quantified (n = 30 and 28 respectively; N=3, *: p<0.05, ***: p<0.001, ****: p<0.0001, One-Way ANOVA, Sídák’s correction). (C) Representative confocal images of the localization of GFPki-Msn and the signal from immunostaining with and anti-Msn antibody (α-Msn). Both channels are displayed separately as inverted greyscale images. Colocalization appears in white in merged images by superposition of the GFPki-Msn signal in green and the anti-Msn signal in magenta. Arrowheads indicates the localization of Msn at the lateral and apical membranes of epithelium follicular cells. Scale bar: 10µm. (D) Representative confocal images show the loss of the Msn staining after expression of a dsRNA against *msn*. The Msn channel is displayed individually as inverted greyscale images. Scale bar: 10µm (E) Representative immunoblotting of HA and actin from lysates of the flies expressing HA-Msn^D160A^, HA-Msn^WT^, HA-Msn^T194A^ used in rescue assay (F) Quantification of HA blot intensities (normalized by actin) of from lysates from flies expressing HA-Msn^D160A^, HA-Msn^WT^, HA-Msn^T194A^ of three independent assays. All error bars represent SEM.

**Figure S3:** Rap2l colocalizes with Msn and affects Msn distribution at the periphery of the cluster without affecting Tao (A) Quantification of *Rap2l* mRNA expression from control and Rap2l-depleted flies in three independent assays (***: p<0.001, One-Way ANOVA, Dunnett’s correction) (B) Representative confocal images of the localization of GFPki-Msn and HA-Rap2l in BCs at the onset. Their colocalization in vesicles is highlighted by red arrowheads in separated channels (shown as inverted greyscale images) and white arrowheads in merged images. Colocalization appears in white in merged images by superposition of the GFPki-Msn signal in green and HA-Rap2l signal in magenta. At the bottom, high magnification image (left) and quantification of the fluorescence signal of Msn and Rap2l by “line scan” (along the yellow, dashed line shown in the high magnification image) (right). Scale bar: 10µm. (C) Quantification of the ratio of Tao-kiVenus mean intensity in the entire cluster, normalized to the signal between nurse cells, in control and Rap2l-depleted clusters. (n = 20, 23 and 23 respectively; N=3, ns= not significant, One-Way ANOVA, Sídák’s correction) (D) Quantification of the enrichment of Tao-kiVenus at inner junctions (BC/BC interface) and the periphery of the cluster (BC/NC interface) at the onset of migration in control and Rap2l-depleted clusters (n = 28, 25 and 28 respectively; N=3, ns= not significant, One-Way ANOVA, Dunnett’s correction) (E) Quantification of the number of Tao-kiVenus positive vesicles per cluster at the onset of migration in control and Rap2l-depleted clusters (n = 28, 27 and 28 respectively; N=3, ns= not significant, One-Way ANOVA, Sídák’s correction) (F) Quantification of *msn* mRNA expression from control and Rap2l-depleted flies. (***: p<0.001, One-Way ANOVA, Dunnett’s correction, N=3) (G) Representative confocal images of the localization of GFPki-Msn at the onset of migration in control and clusters overexpressing HA-Rap2l, displayed as inverted greyscale images. Scale bar: 10µm. (H) Quantification of the number of GFPki-Msn positive vesicles per cluster at the onset of migration in control and clusters overexpressing HA-Rap2l. (n = 23 and 28 respectively; n represents the number of independent BC clusters; N=3 experiments, ns= not significant, Mann-Whitney test). (I) Quantification of the ratio of GFPki-Msn mean intensity in the entire cluster, normalized to the signal between nurse cells, in control and clusters overexpressing HA-Rap2l. (n = 20 and 24 respectively; N=3, ns= not significant, Mann-Whitney test). (J) Quantification of the enrichment of GFPki-Msn at inner junctions (BC/BC interface) and at the periphery of the cluster (BC/NC interface) at the onset of migration in control and clusters overexpressing HA-Rap2l. (n = 23 and 28 respectively; N=3, ns= not significant, ***: p<0.001, ****: p<0.0001, One-Way ANOVA, Sídák’s correction). All error bars represent SEM.

**Figure S4:** A subset of Msn localizes in early endosomes. (A) Representative confocal images showing co-labelling of Msn by immunostaining and early endosomes (marked with GFP-Rab5) in BCs. The localization in vesicles is highlighted by red arrowheads in separated channels (shown as inverted greyscale images) and by white arrowheads in merged images. On the right, high magnification image. Scale bar: 10μm (B) Quantification of the fluorescence signal of Msn and Rab5 by “line scan” (along the yellow, dashed line shown in A) (C) Representative confocal images showing co-labelling of Msn by immunostaining and recycling endosomes (marked with GFP-Sec15) in BC clusters. Each channel is presented separated as inverted greyscale images) and merged with Sec15 in green and Msn in magenta. No obvious overlap is observed. Scale bar: 10μm.

