## Supplementary figures and images for "Dual regulation of Misshapen by Tao and Rap2l promotes collective cell migration"

### Supplemental figures

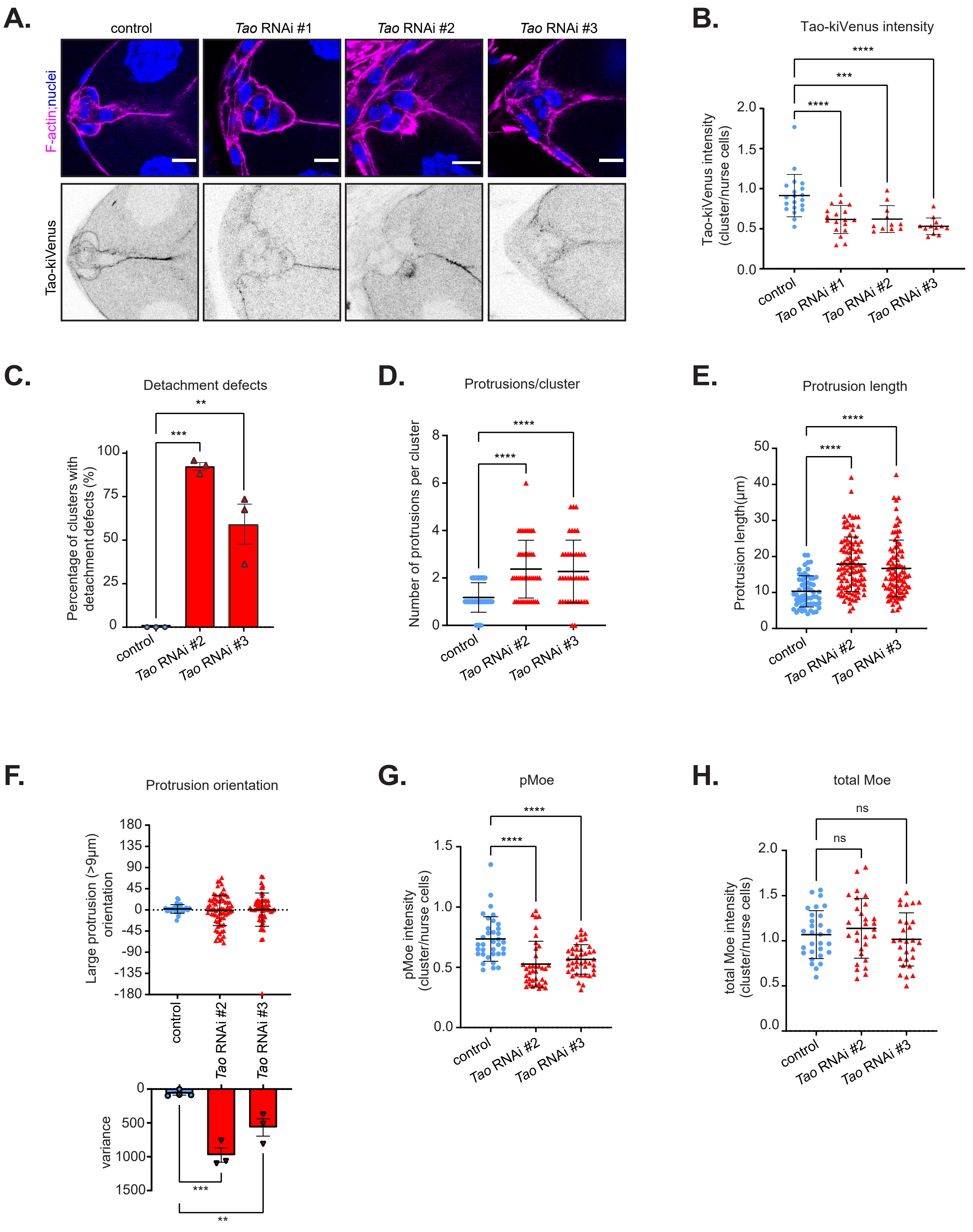

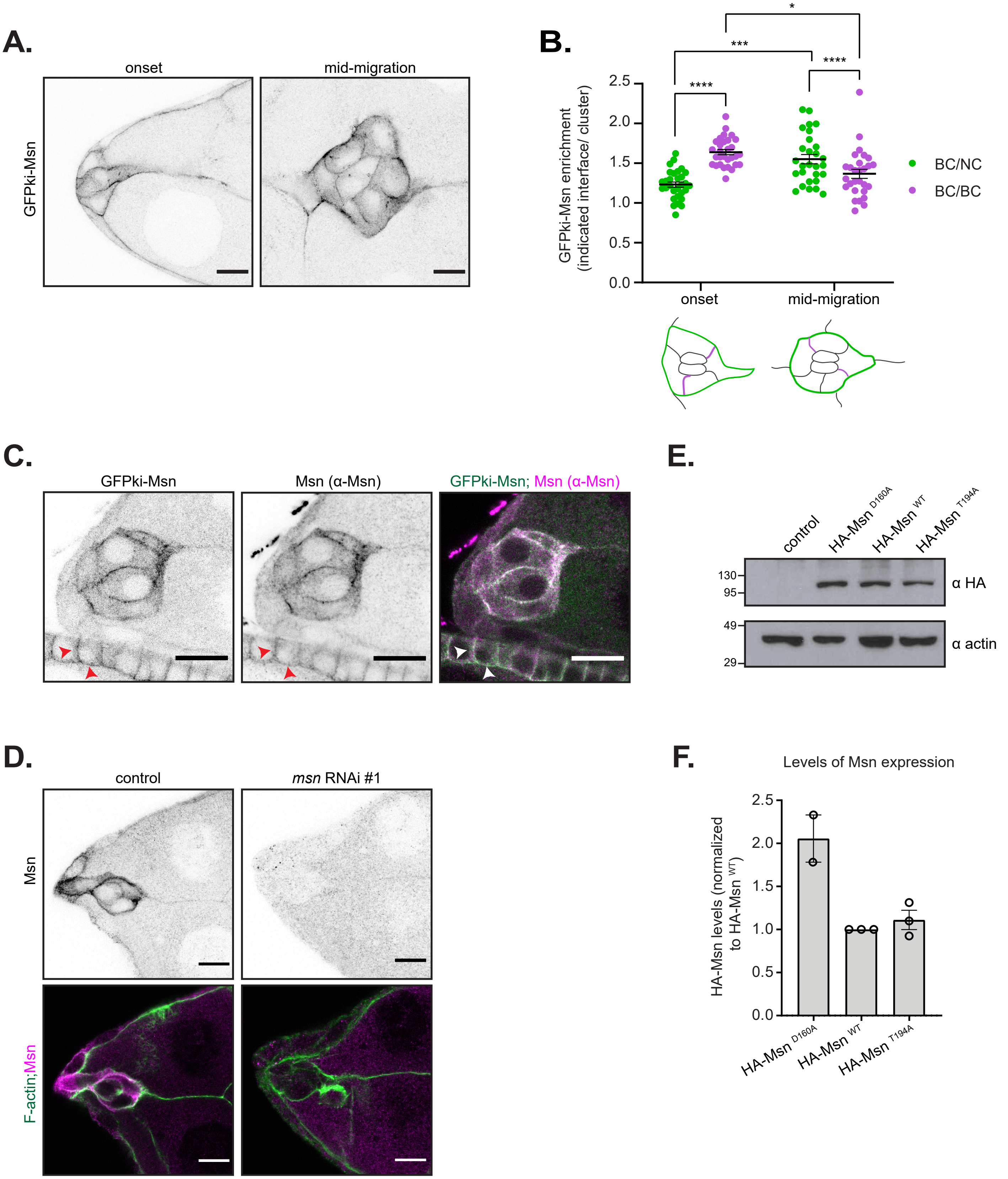

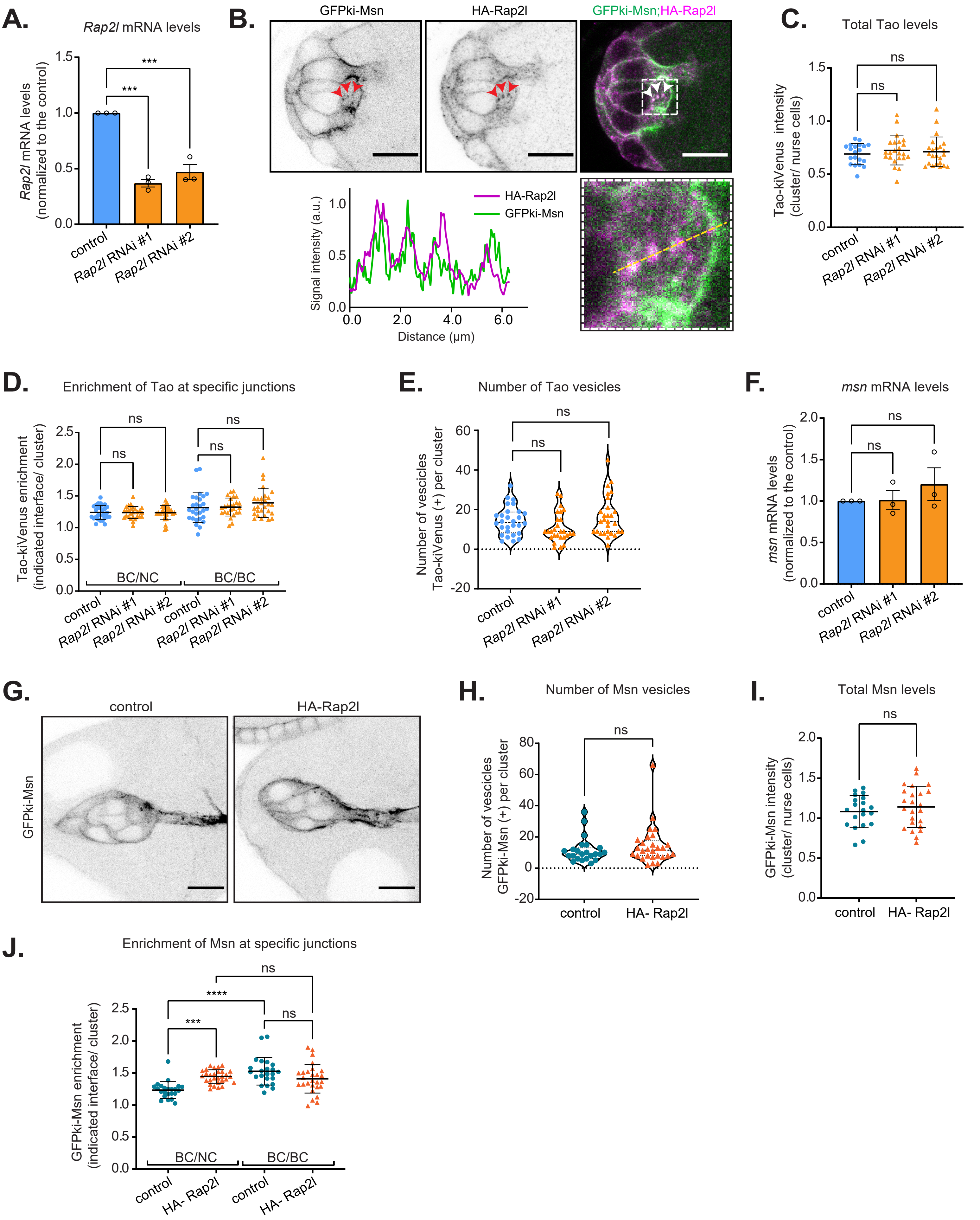

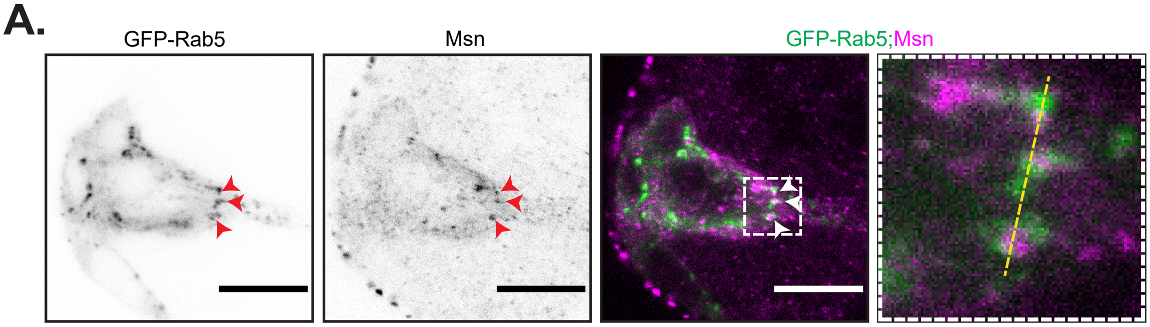

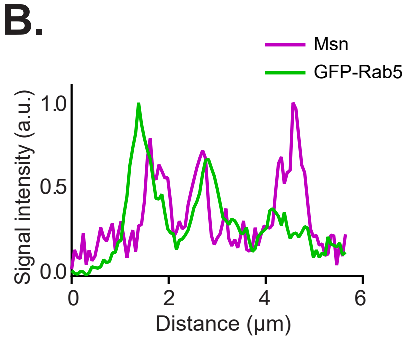

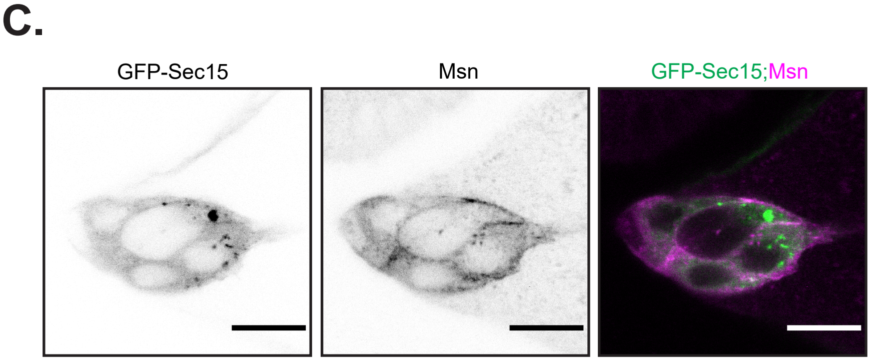
